# Uncovering novel mutational signatures by *de novo* extraction with SigProfilerExtractor

**DOI:** 10.1101/2020.12.13.422570

**Authors:** S M Ashiqul Islam, Yang Wu, Marcos Díaz-Gay, Erik N Bergstrom, Yudou He, Mark Barnes, Mike Vella, Jingwei Wang, Jon W Teague, Peter Clapham, Sarah Moody, Sergey Senkin, Yun Rose Li, Laura Riva, Tongwu Zhang, Andreas J Gruber, Raviteja Vangara, Christopher D Steele, Burçak Otlu, Azhar Khandekar, Ammal Abbasi, Laura Humphreys, Natalia Syulyukina, Samuel W Brady, Boian S Alexandrov, Nischalan Pillay, Jinghui Zhang, David J Adams, Iñigo Martincorena, David C Wedge, Maria Teresa Landi, Paul Brennan, Michael R Stratton, Steven G Rozen, Ludmil B Alexandrov

## Abstract

Mutational signature analysis is commonly performed in genomic studies surveying cancer and normal somatic tissues. Here we present SigProfilerExtractor, an automated tool for accurate *de novo* extraction of mutational signatures for all types of somatic mutations. Benchmarking with a total of 33 distinct scenarios encompassing 1,900 simulated signatures operative in more than 60,000 unique synthetic genomes demonstrates that SigProfilerExtractor outperforms thirteen other tools across all datasets with and without noise. For simulations with 5% noise, reflecting high-quality genomic datasets, SigProfilerExtractor outperforms other approaches by elucidating between 20% and 50% more true positive signatures while yielding more than 5-fold less false positive signatures. Applying SigProfilerExtractor to 2,778 whole-genome sequenced cancers reveals three previously missed mutational signatures. Two of the signatures are confirmed in independent cohorts with one of these signatures associating with tobacco smoking. In summary, this report provides a reference tool for analysis of mutational signatures, a comprehensive benchmarking of bioinformatics tools for extracting mutational signatures, and several novel mutational signatures including a signature putatively attributed to direct tobacco smoking mutagenesis in bladder cancer and in normal bladder epithelium.

## INTRODUCTION

*De novo* extraction of mutational signatures^1^ is an unsupervised machine learning approach where a matrix, ***M***, which corresponds to the somatic mutations in a set of cancer samples under a mutational classification^2^, is approximated by the product of two low-rank matrices, ***S*** and ***A***. The matrix ***S*** reflects the set of mutational signatures while the matrix ***A*** encompasses the activities of the signatures; an activity corresponds to the number of mutations contributed by a signature in a cancer sample. Algorithmically, *de novo* extraction of mutational signatures has relied on nonnegative matrix factorization (NMF)^3^ or on approaches mathematically analogous to NMF^4–6^. The main advantage of NMF over other factorization approaches is its ability to yield nonnegative factors that are part of the original data, thus, allowing interpretation of the identified nonnegative factors^3^. Biologically, mutational signatures extracted from cancer genomes have been attributed to exposures to environmental carcinogens, failure of DNA repair pathways, infidelity/deficiency of replicating polymerases, iatrogenic events, and others^7–14^.

Since we introduced the mathematical concept of mutational signatures^1^, a number of computational frameworks have been developed for performing *de novo* extraction of mutational signatures (**Table 1**)^14–27^. Notably, the majority of existing *de novo* extraction tools *(i)* predominately support the simplest mutational classification, *viz.*, SBS-96 which encompasses single base substitutions with their immediate 5’ and 3’ sequence context^2^; *(ii)* lack automatic selection for the number of mutational signatures; *(iii)* do not identify a robust solution leading to different results following re-analysis of the same dataset; *(iv)* require pre-selection of a large number of priors and/or hyperparameters; *(v)* do not decompose *de novo* signatures to the set of reference COSMIC signatures^14^. Importantly, there has been no extensive benchmark of the existing tools for *de novo* extraction leading to uncertainty regarding their performance.

**Table 1:**
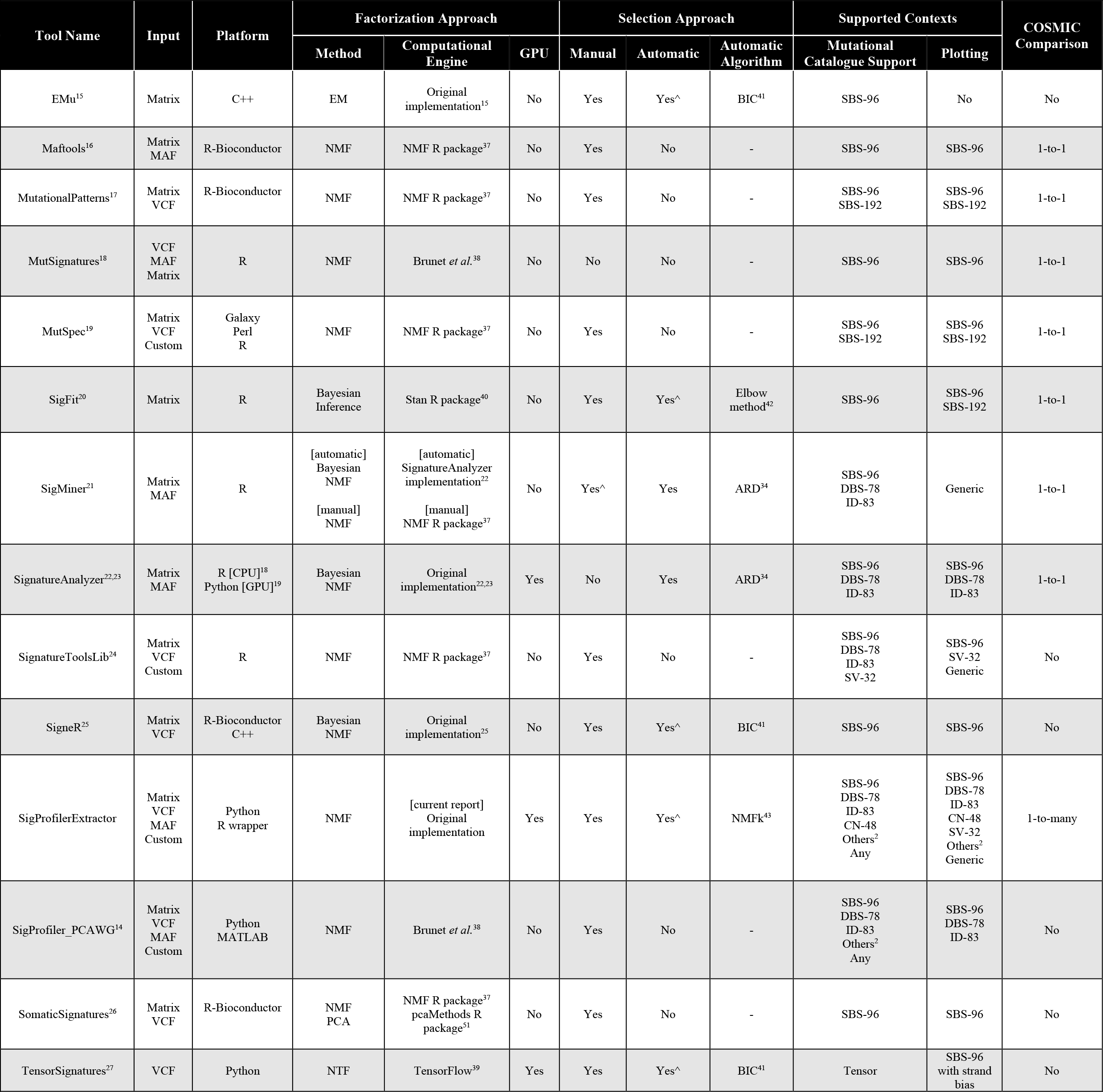
Overview of bioinformatics tools for *de novo* extraction of mutational signatures. Tools are ordered alphabetically. Notations: ^ denotes the default approach for selecting the total number of signatures when a tool supports both manual and automatic selection; 1-to-1 refers to one *de novo* signature being matched with exactly one COSMIC signature; 1-to-many refers to one *de novo* signature being matched with a combination of one or more COSMIC signatures. Abbreviations: MAF: mutation annotation format; VCF: variant call format; EM: expectation-maximization algorithm; NMF: nonnegative matrix factorization; NTF: nonnegative tensor factorization; ARD: automatic relevance determination; BIC: Bayesian information criterion; COSMIC: catalogue of somatic mutations in cancer; SBS: single base substitutions; DBS: doublet base substitutions; ID: small insertions and deletions; SV: structural variants.

To address these limitations, here, we present SigProfilerExtractor – a reference tool for *de novo* extraction of mutational signatures. SigProfilerExtractor allows analysis of all types of mutational classifications, performs automatic selection of the number of signatures, yields robust solutions, requires only minimum setup, and decomposes *de novo* extracted signatures to known COSMIC signatures. A comprehensive benchmark including 3,448 unique matrix decompositions with SigProfilerExtractor and thirteen other tools across a total of 33 distinct scenarios reveals that SigProfilerExtractor is robust to noise and that it outperforms all other computational tools for *de novo* extraction of mutational signatures (**Supplementary Tables 1–3**). Applying SigProfilerExtractor to the recently published set of 2,778 whole-genome sequenced cancers from the Pan-Cancer Analysis of Whole Genomes (PCAWG) project^28^ elucidates three novel signatures that were not found in the original PCAWG analysis of mutational signatures^14^. Two of the signatures are confirmed in independent cohorts and a putative etiology of tobacco-associated mutagenesis is attributed to one of these signatures.

## RESULTS

### Overview of SigProfilerExtractor and its implementation

SigProfilerExtractor is implemented as a Python package, with an R wrapper, allowing users to run it in both Python and R environments: https://github.com/AlexandrovLab/SigProfilerExtractor. The tool is also extensively documented including a detailed Wiki page: https://osf.io/t6j7u/wiki/home/. By default, the tool requires only a single parameter – the input dataset containing the mutational catalogues of interest. SigProfilerExtractor supports most used formats outputted by variant calling algorithms (*e.g.*, VCF, MAF, *etc.*), which are internally converted to a matrix, ***M***, by SigProfilerMatrixGenerator^2^. SigProfilerExtractor can also be applied to a text file containing a matrix, ***M***, thus supporting nonnegative matrix factorization for any custom matrix dataset. By default, the tool decomposes the matrix ***M*** searching for an optimal solution for the number of operative signatures, ***k***, between 1 and 40 mutational signatures (**Figure 1*a***). For each decomposition, SigProfilerExtractor performs 500 independent factorizations and, for each repetition, the matrix ***M*** is first Poisson resampled and normalized and, subsequently, factorized with the multiplicative update NMF algorithm^3^ by minimizing an objective function based on the Kullback–Leibler divergence measure^29^ (**Figure 1*b***). Custom partition clustering, that utilizes the Hungarian algorithm^30^ for comparing different repetitions, is applied to the 500 factorizations to identify stable solutions^31^ (**Figure 1*b***). Specifically, SigProfilerExtractor selects the centroids of stable clusters as optimal solutions, thus, making these solutions resistant to fluctuations in the input data and to the lack of uniqueness of NMF due to the potential existence of multiple convergent stationary points in the solution^32^. Lastly, when applicable, the optimal set of *de novo* signatures are matched to the set of reference COSMIC mutational signatures (**Figure 1*c***) with any *de novo* signature reported as novel when it cannot be decomposed by a combination of known COSMIC signatures.

**Figure 1.**
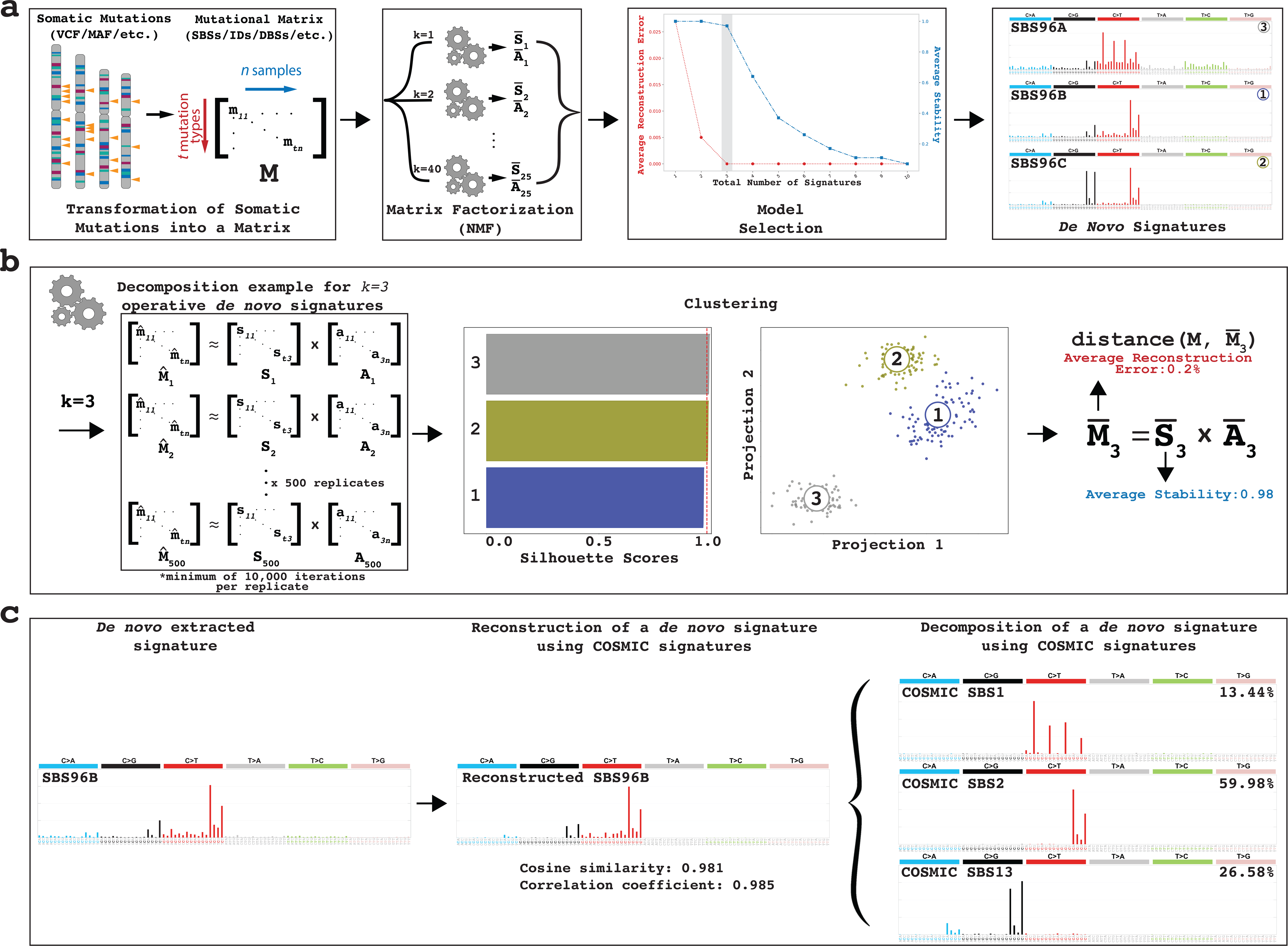
Overview of SigProfilerExtractor. ***(a)*** SigProfilerExtractor’s general workflow is outlined starting from an input of somatic mutations and resulting in an output of *de novo* mutational signatures. An example is shown for a solution with three *de novo* signatures. Somatic mutations are first converted into a mutational matrix. Subsequently, the matrix is factorized with different ranks using nonnegative matrix factorization. Model selection is applied to identify the optimal factorization rank based on each solution’s stability and its reconstruction of the original data. ***(b)*** Schematic representation for an example decomposition with a factorization rank of *k=3* reflecting three operative mutational signatures. By default, SigProfilerExtractor performs 500 independent nonnegative matrix factorizations with the matrix ***M*** being Poisson resampled and normalized (denoted by ^) prior to each factorization. Partition clustering of the 500 factorizations is used to evaluate the factorization stability rank, measured in silhouette values; clustering can also be presented as two-dimensional projections revealing more similar mutational signatures as shown for the three example signatures. The centroid of the clustered solutions (denoted by **–**) is compared to the original matrix ***M***. ***(c)*** All identified *de novo* signatures are matched to a combination of known COSMIC mutational signatures. An example is given for *de novo* extracted signature SBS96B which matches a combination of COSMIC signatures SBS1, SBS2, and SBS13.

### Framework for benchmarking tools for de novo extraction of mutational signatures

To allow comprehensive benchmarking of tools for *de novo* extraction of mutational signatures, more than 60,000 unique synthetic cancer genomes were generated with known ground-truth mutational signatures (**Supplementary Note 1**). These synthetic data included 32 distinct noiseless scenarios and one scenario with five different levels of noise. Each scenario contained between 3 and 40 known signatures operative in 200 to 3,000 simulated cancer genomes (**Supplementary Tables 1–3**). Some of the scenarios were generated up to 20 times to account for variability in the simulated data. Most noiseless scenarios (20/32) were based on SBS-96 mutational classification; 12 scenarios based on extended mutational classifications, *i.e.,* matrices with more than 96 mutational channels, were also included (**Supplementary Table 3**). To avoid bias in evaluating each tool’s performance, three sets of SBS-96 mutational signatures were used for generating the synthetic data: *(i)* COSMICv3 reference signatures^14^; *(ii)* SA signatures previously extracted by SignatureAnalyzer^14^; and *(iii)* randomly generated signatures. For presentation simplicity, scenarios were labeled based on their complexity as easy, medium, or hard. Easy scenarios were generated using ≤5 signatures and provide a good indication of each tool’s performance on approximately 7.4% of human cancer types (*e.g.*, pediatric brain tumors). Medium scenarios contained 11 to 21 signatures and biologically reflect 15.9% of cancer types (*e.g.*, cervical cancer). Hard scenarios have more than 25 signatures and reflect 59.5% of human cancer types (*e.g.*, breast, lung, liver, *etc.*) as well as pan-cancer datasets. In addition to the 32 noiseless scenarios, one SBS-96 scenario with five different levels of noise, ranging between 0% and 10%, was included in the benchmark (**Supplementary Note 1**).

To compare the performance between different tools for *de novo* extraction of mutational signatures, we developed a standard set of evaluation metrics (**Supplementary Figure 1**). Specifically, each *de novo* extracted signature is classified as either a *true positive* (TP), *false positive* (FP), or *false negative* (FN) signature. An extracted signature is considered TP if it matches one of the ground-truth signatures above a cosine similarity threshold of 0.90. In contrast, a signature is classified as FP when it has a maximum cosine similarity below 0.90 with all ground-truth signatures. Lastly, FN signatures are ground-truth signatures that were not detected in the data. These standard metrics allow calculating each tool’s precision, sensitivity, and F_1_ score. Precision is defined as 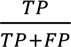, sensitivity as 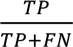, and F_1_ score is the harmonic mean of the precision and sensitivity: 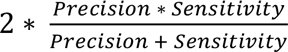.

### Benchmarking SigProfilerExtractor and thirteen other tools using SBS-96 noiseless data

SigProfilerExtractor and thirteen other tools (**Table 1**) were first applied to all noiseless scenarios based on the SBS-96 mutational classification. The thirteen tools include SignatureAnalyzer and SigProfiler_PCAWG, a legacy MATLAB/Python version of SigProfilerExtractor, which were jointly used in the PCAWG analysis of mutational signatures and the derivation of the COSMICv3 set of reference mutational signatures^14^. Except for MutSignatures which can only decompose a matrix for a fix number of signatures, all other tools were applied to each scenario by using their suggested methods for selecting the number of operative signatures. Apart from SignatureAnalyzer which lacks this capability, all other tools were forced to extract the known number of ground-truth signatures. Results from the suggested approach reflect the expected outcome from running a tool on an unknown dataset, while results from the forced approach allow understanding limitations in each tool’s implementation. Our evaluation reveals that most tools can successfully extract mutational signatures from easy scenarios with the majority of F_1_ scores between 0.90 and 1.00 (**Figure 2*a***). This is perhaps unsurprising as many of these tools used synthetic data with ≤5 signatures to evaluate their performance in their respective original publications^15–17, 19–26^. In contrast, medium scenarios have proven to be a challenge for most tools with only SigProfilerExtractor, SigProfiler_PCAWG, and SignatureAnalyzer exhibiting F_1_ scores above 0.90. All tools had worst performance for the hard set of scenarios with F_1_ scores below 0.80; only SigProfilerExtractor had an F_1_ score of almost 0.90 (**Figure 2*a***).

**Figure 2.**
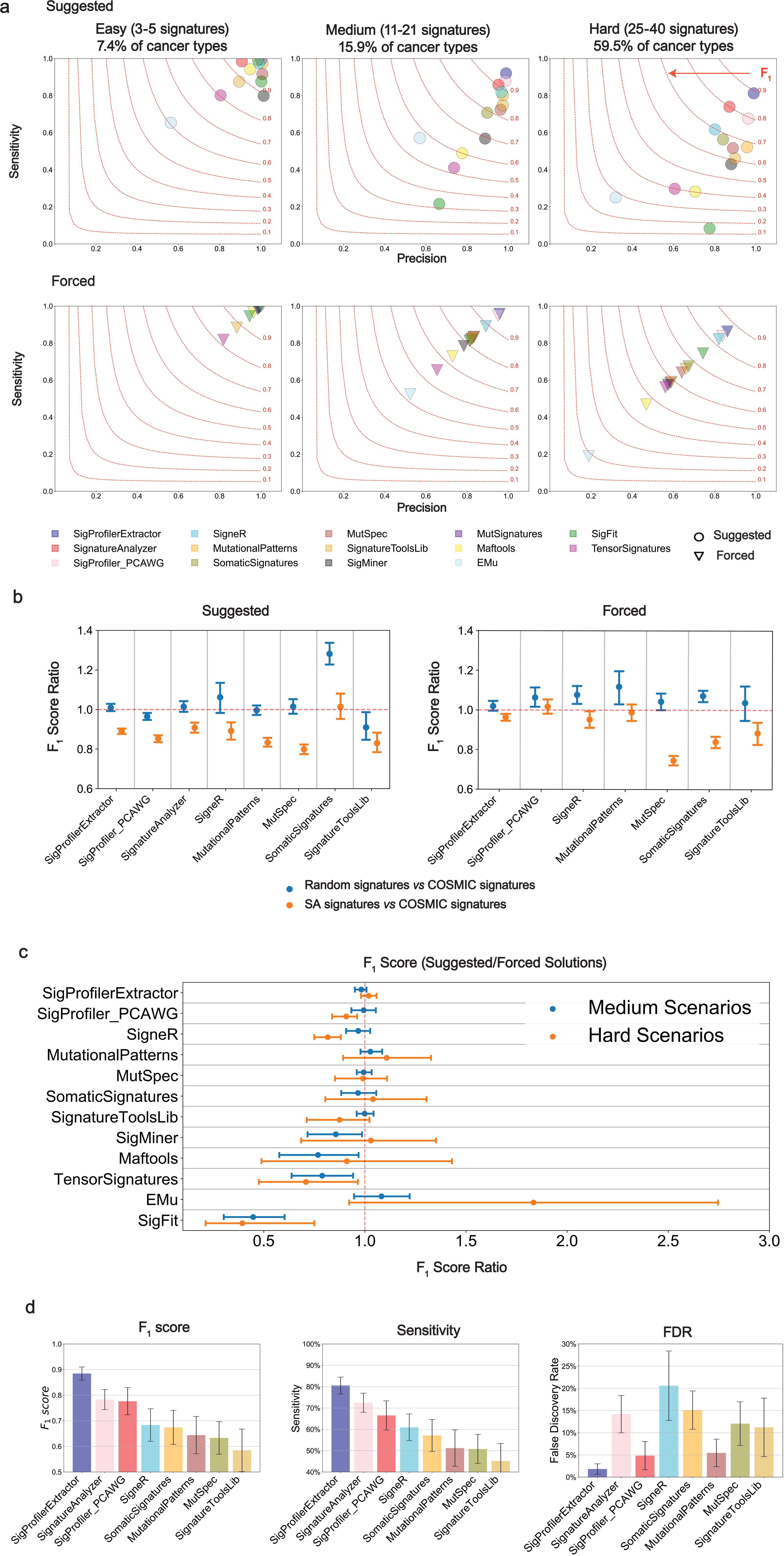
Benchmarking of bioinformatics tools for *de novo* extraction of mutational signatures using SBS-96 noiseless scenarios. ***(a)*** Average precision (x-axes), sensitivities (y-axes), and F_1_ scores (harmonic mean of precision and sensitivity; red curves) are shown across the three types of scenarios. Different tools are displayed using circles and triangles with different colors. Circles are used to display results for suggested model selection, which most closely matches analysis of a real dataset. Triangles are used to display results for forced model selection, where tools were required to extract the known total number of ground-truth mutational signatures. All triangles are located on the diagonal as the forced model selection results in equal numbers of false positive and false negative signatures. ***(b)*** Evaluating the effect of ground-truth signatures on the *de novo* extraction by different tools (x-axes). Ratio of F_1_ scores (y-axes) with confidence intervals were calculated for medium complexity scenarios simulated using COSMIC, SA, or random signatures. Ratio of approximately 1.00 indicates a similar performance between different types of signatures. ***(c)*** Evaluating the performance of *de novo* extraction between suggested and forced selection for different tools (x-axes). Ratio of F_1_ scores (y-axes) with confidence intervals were calculated for all medium and hard scenarios. Ratio of approximately 1.00 indicates a similar performance between suggested and forced model selection. ***(d)*** Summary of the performance for the top seven tools on hard SBS-96 noiseless scenarios with suggested model selection. Y-axes reflect F_1_ score (left plot), sensitivity (middle plot), and false discovery rate (right plot), respectively. Results from SignatureAnalyzer and MutSignatures are not displayed in panels ***(a)***, ***(b)***, and ***(c)*** for forced and suggested model selections, respectively, as the tools do not support these types of analyses.

To evaluate whether the type of ground-truth signatures affects the *de novo* extraction, we compared the ratio of F_1_ scores (rF_1_) from scenarios generated using COSMIC, SA, or random signatures (**Figure 2*b***). Most tools had similar performance (rF_1_≈1) between COSMIC and random signatures and worst performance with SA signatures (rF_1_<1). SomaticSignatures was an exception as it performed well on random signatures but had similarly suboptimal performance on COSMIC and SA signatures. SigProfilerExtractor outperformed all other tools regardless of whether the synthetic data were generated using COSMIC, SA, or random signatures (**Supplementary Table 1**).

To examine the performance of *de novo* extraction between the suggested and forced selection of the total number of signatures, we evaluated rF_1_ across all medium and hard scenarios (**Figure 2*c***). SigProfilerExtractor exhibited almost identical F_1_ scores between the suggested and forced selection indicating a good performance of the automatic selection algorithm. Most other tools had similar F_1_ scores between the suggested and forced selection albeit with more variability across the different scenarios (**Figure 2*c***). For example, MutSpec had rF_1_≈1 in both medium and hard scenarios indicating that MutSpec is performing worse than SigProfilerExtractor (**Figure 2*a***) not because of its algorithm for selecting the total number of signatures but likely due to its implementation of the utilized numerical factorization. SigneR (hard scenarios), SigProfiler_PCAWG (hard), SigMiner (medium), TensorSignatures (all), and SigFit (all) had lower F_1_ scores for automatic solutions compared to forced solutions (rF_1_<1), thus, indicating that their automatic approaches for selecting the total number of signatures are not optimally performing (**Figure 2*c***). Surprisingly, EMu had higher F_1_ scores for automatic solutions in some hard scenarios. Considering the overall performance of EMu (**Figure 2*a***), this outcome likely reflects the lack of convergence during the minimization of the EMu objective function for certain number of signatures in the hard scenarios.

Overall, across all suggested extractions from noiseless hard scenarios reflecting ∼60% of human cancer types, SigProfilerExtractor outperformed all other tools. SigProfilerExtractor was able to identify between 10% and 37% more true positive signatures while yielding between 2.7- and 16-fold less false positive signatures compared to the next seven best performing tools: SigProfiler_PCAWG, SignatureAnalyzer, SigneR, MutationalPatterns, MutSpec, SomaticSignatures, and SignatureTools (**Figure 2*d*** and **Supplementary Table 1**).

### Extended benchmarking of SigProfilerExtractor and the other seven top performing tools

The reported comparisons for SBS-96 scenarios rely on a cosine similarity ≥0.90 for determining TP signatures and <0.90 for determining FP signatures. Note that a cosine similarity ≥0.90 is highly unlikely to happen purely by chance (p-value = 5.90 x 10^-9^) as two random nonnegative vectors are expected to have an average cosine similarity of 0.75 purely by chance^33^. Importantly, SigProfilerExtractor’s performance does not depend on the specific value of the cosine similarity threshold (**Figure 3*a***) as the tool consistently outperforms other bioinformatics approaches for almost any value of the threshold above 0.80 (p-value: 0.057). Cosine similarity thresholds below 0.80 were not explored as extracted signature may be similar to ground-truth signatures purely by chance.

**Figure 3.**
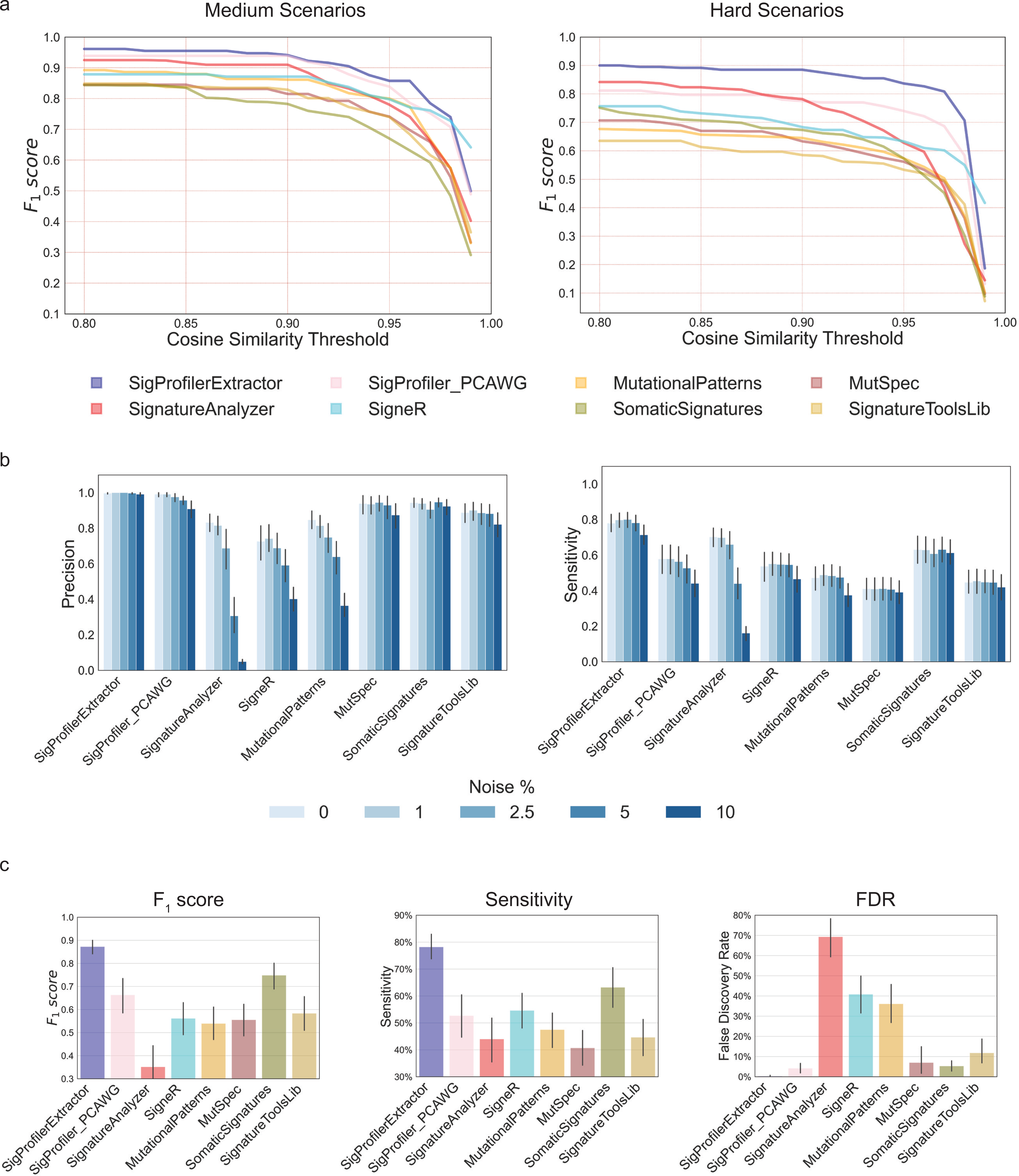
Additional evaluations of the top seven bioinformatics tools for *de novo* extraction of mutational signatures. ***(a)*** Average F_1_ scores for the top seven tools based on different thresholds for cosine similarity in suggested medium and hard scenarios; thresholds for cosine similarity are used for determining true positive signatures (**Supplementary** Figure 1). X-axes reflect the cosine similarity thresholds, while the Y-axes correspond to the average F_1_ scores corresponding to cosine similarity thresholds. ***(b)*** Precision and sensitivity of the top seven tools for SBS-96 scenarios with different levels of noise. Noise levels reflect the average number of somatic mutations in a cancer genome affected by additive white Gaussian noise; for example, 1% noise corresponds to approximately 1% of mutations in a sample being due to noise. ***(c)*** Summary of the performance of the top seven tools on SBS-96 scenarios with 5% noise. Y-axes reflect F_1_ score (left plot), sensitivity (middle plot), and false discovery rate (right plot), respectively.

Additional benchmarking was performed by generating 12 scenarios simulated using between 3 and 30 signatures with an extended number of mutational channels (**Supplementary Note 1**). SigProfilerExtractor and SignatureAnalyzer are the only two tools that support analysis of custom size matrices and provide GPU support (**Table 1**), thus, allowing analysis of data with extended number of mutational channels within a reasonable timeframe. In contrast, all other matrix factorization tools rely solely on CPU implementations with full runs expected to take many months for each tool applied to these scenarios (**Table 1**). SigProfilerExtractor and SignatureAnalyzer exhibited similar performance on the extended noiseless scenarios to that observed on SBS-96 noiseless scenarios. Overall, SigProfilerExtractor outperformed SignatureAnalyzer with average F1 scores of 0.92 and 0.85, respectively (**Supplementary Table 2**).

To further compare SigProfilerExtractor with the other seven top performing tools, we applied each tool to a dataset with 30 ground-truth SBS-96 signatures operative in 1,000 genomes and random noise between 0% and 10%. Analysis for each noise level was repeated 20 times to account for any variability in the noise generation. SigProfilerExtractor, SomaticSignatures, MutSpec, SignatureToolsLib were robust to noise with mostly unaffected performance (**Figure 3*b*** and **Supplementary Table 3**). In contrast, SigProfiler_PCAWG, SignatureAnalyzer, SigneR, and MutationalPatterns were susceptible to noise (**Figure 3*b***). For example, 2.5% noise reduced SignatureAnalyzer’s F_1_ from 0.76 to 0.66 while 10% noise reduced its F_1_ to 0.07. Similarly, 10% noise reduced the F_1_ of SigProfiler_PCAWG from 0.76 to 0.57, the F_1_ of SigneR from 0.61 to 0.43, and the F_1_ of MutationalPatterns from 0.60 to 0.37. SignatureAnalyzer’s reduced performance on data with noise is due to its automated approach for selecting total number of signatures. SignatureAnalyzer uses automatic relevance determination^34^ for selecting the number of signatures with this number increasing from 26 (no noise; 30 ground-truth signatures) to 96 signatures (10% noise; **Supplementary Table 3**). In contrast, SigProfiler_PCAWG, SigneR and MutationalPatterns exhibit similar performance between forced and suggested solutions on data with noise (**Supplementary Table 3**) indicating that their reduced performance is likely due to the numerical implementation of their respective factorization approaches.

SigProfilerExtractor outperformed all other tools regardless of the levels of noise. Simulations with 5% noise reflect genomics datasets with ∼0.95 average sensitivity and precision of single base substitutions, similar to the recently published PCAWG cohort which has 95% sensitivity (90% confidence interval, 88–98%) and 95% precision (90% confidence interval, 71–99%)^28^. For simulations with 5% noise, SigProfilerExtractor was able to identify between 20% and 50% more true positive signatures while yielding more than 5-fold less false positive signatures compared to the next seven best performing tools: SigProfiler_PCAWG, SignatureAnalyzer, SigneR, MutationalPatterns, MutSpec, SomaticSignatures, and SignatureTools (**Figure 3*c*** and **Supplementary Table 3**).

### Analysis of 2,778 whole-genome sequenced human cancers with SigProfilerExtractor

To demonstrate its ability to yield novel biological results, SigProfilerExtractor was applied to the recently published set of 2,778 whole-genome sequenced cancers^28^. As previously done in our original PCAWG analysis of mutational signatures^14^, extraction of mutational signatures was performed within each cancer type as well as across all samples (**Supplementary Data**). In addition to all previously detected signatures^14^, our direct application of SigProfilerExtractor revealed three novel mutational signatures were identified in the PCAWG dataset: SBS92, SBS93, and SBS94 (**Figure 4** and **Supplementary Table 4**).

**Figure 4.**
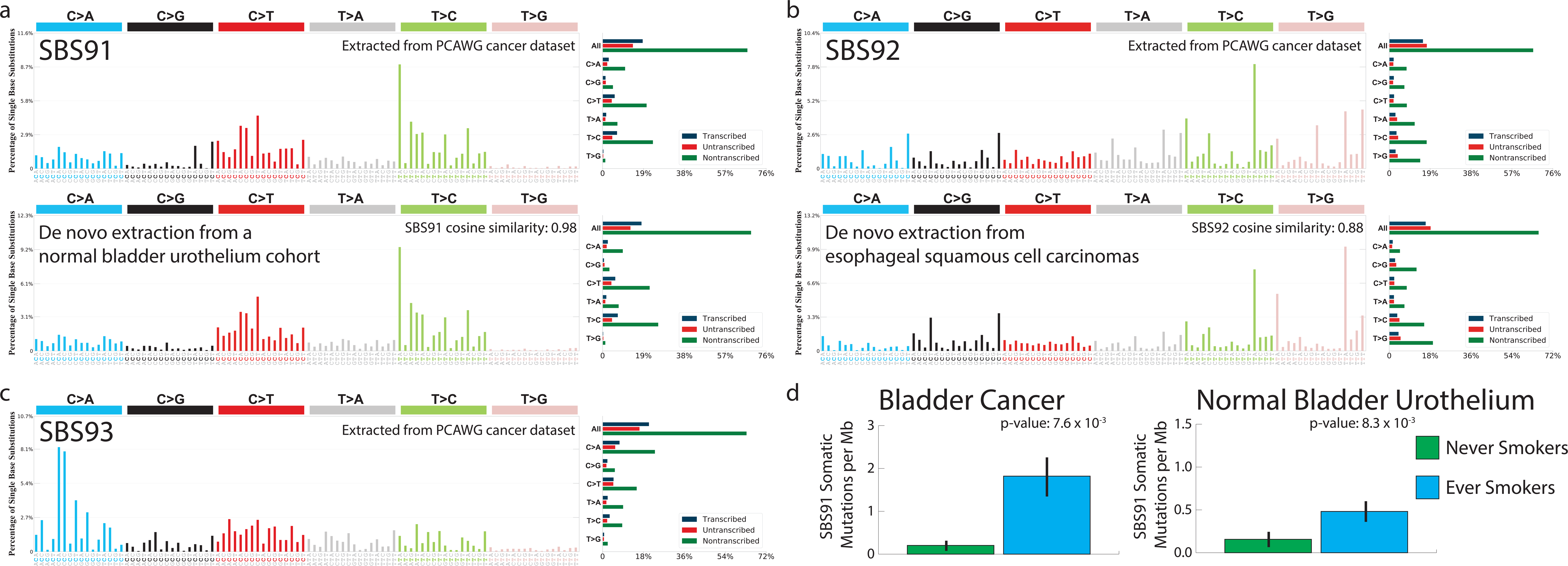
Novel signatures identified in the PCAWG cohort of 2,778 whole-genome sequenced cancers. Mutational signatures are displayed using 96-plots. Single base substitutions are shown using the six subtypes of substitutions: C>A, C>G, C>T, T>A, T>C, and T>G. Underneath each subtype are 16 bars reflecting the sequence contexts determined by the four possible bases 5’ and 3’ to each mutated base. Additional information whether mutations from a signature are in non-transcribed/intergenic DNA, on the transcribed strand of a gene, or on the untranscribed strand of the gene is provided adjacent to the 96 plots. ***(a)*** Mutational profile of signature SBS92 derived from the PCAWG cohort (top). Confirmation of the profile of signature SBS92 (bottom) by analysis of an independent whole-genome sequenced set of microbiopsies of histologically normal urothelium^35^. ***(b)*** Mutational profile of signature SBS93 derived from the PCAWG cohort (top). Confirmation of the profile of signature SBS93 (bottom) by analysis of an independent whole-genome sequenced set of esophageal squamous cell carcinomas^28^. ***(c)*** Mutational profile of signature SBS94 derived from the PCAWG cohort. Signature SBS94 was not identified in any additional independent cohort. ***(d)*** Bars are used to display average values for numbers of somatic substitutions per megabase (Mb) attributed to signature SBS92 in bladder cancer and normal bladder urothelium. Green bars represent never-smokers, whereas blue bars correspond to ever smokers. Error bars correspond to 95% confidence intervals. Each p-value is based on a Wilcoxon rank sum test.

Signature SBS92 was found predominately in PCAWG bladder cancers; the signature was characterized by T>C mutations with strong transcriptional strand bias consistent with damage on purines for all types of single base substitutions (**Figure 4*a***). Signature SBS92 was 9-fold elevated (**Figure 4*d***; p-value: 7.6 x 10^-3^ using Wilcoxon rank sum test) in bladder cancers of ever smokers compared to never-smokers in the PCAWG cohort. An almost identical signature was identified by re-analyzing a recently published cohort of 88 whole-genome sequenced microbiopsies of histologically normal urothelium^35^ with the similarity extending to both trinucleotide context and transcriptional strand bias (**Figure 4*a***; cosine similarity: 0.98; p-value < 10^-256^). Consistently, SBS92 was found to be 3-fold elevated in the normal urothelium of tobacco ever smokers compared to never-smokers (**Figure 4*d***; p-value: 8.3 x 10^-3^ using Wilcoxon rank sum test).

Signature SBS93 was identified almost exclusively in PCAWG stomach cancers. SBS93 was characterized by T>C and T>G mutations with a strand bias consistent with damage on pyrimidines for Tp<u>T</u>pA contexts (mutated base underlined; **Figure 4*b***). *De novo* extraction from the Mutographs cohort of 552 whole-genome sequenced esophageal squamous cell carcinomas^36^, a cancer type not included in the PCAWG dataset^28^, identified an analogous mutational signature with the similarity extending to both trinucleotide context and transcriptional strand bias (**Figure 4*b***; cosine similarity: 0.88; p-value: 1.1 x 10^-6^). Signature SBS94 was found at high levels in a single colorectal PCAWG cancer with smaller contributions to another 8 colorectal cancers. The pattern of SBS94 was characterized by C>A mutations with a strand bias indicative of damage on guanine (**Figure 4*c***). Validation of somatic mutations by visual inspection confirmed that 98% of mutations contributed by SBS94 are likely real. Signatures SBS93 and SBS94 did not associate with any of the available PCAWG metadata^28^ and their etiologies remain unknown.

## DISCUSSION

The performed large-scale benchmarking demonstrates that SigProfilerExtractor outperforms thirteen other tools for *de novo* extraction of mutational signatures for noiseless datasets as well as for datasets containing matrices with different levels of random noise. Importantly, SigProfilerExtractor generates almost no false positive signatures while still identifying a higher number of true positive signatures when compared to any of the other tools (**Figure 2*d*** and **Figure 3*c***). *De novo* extraction of mutational signatures relies both on a factorization approach and on a model selection algorithm for determining the total number of operative signatures (**Figure 1**). Benchmarking with forced model selection, where tools were required to extract the known number of ground-truth mutational signatures, reveals that SigProfilerExtractor’s factorization performs better when compared to the factorizations of other tools (**Figure 2*a*** and **Supplementary Tables 1-3**). Similarly, benchmarking with suggested model selection, which most closely matches analysis of a real dataset with unknown number of signatures, further demonstrates SigProfilerExtractor’s ability to reveal novel biological results (**Figure 2*a*** and **Supplementary Tables 1-3**).

While our benchmarking evaluated thirteen additional tools, six of the thirteen tools internally rely on the same computational engine. Maftools, MutationalPatterns, MutSpec, SignatureToolsLib, SigMiner, and SomaticSignatures use the NMF R package^37^ to perform their factorization (**Table 1**), albeit with slightly different hyperparameters and, in some cases, distinct pre-processing of the input matrix. Predictably, these six tools have similar performance across many of the scenarios (**Supplementary Tables 1-3**). SigProfiler_PCAWG and MutSignatures utilize customize versions of an NMF implementation originally developed by Brunet *et al.*^38^ for analysis of gene expression data. TensorSignatures makes use of the standard factorization algorithms included in TensorFlow^39^. SigFit uses a previously developed nonnegative factorization method, *viz.*, Stan R package^40^. In contrast, EMu, SignatureAnalyzer, SigneR, and SigProfilerExtractor provide original implementations of their factorization algorithms (**Table 1**). EMu was originally developed and tested on small datasets (*e.g.,* 21 breast genomes)^15^ and its benchmarking performance is perhaps unsurprising considering the large number of synthetic samples used in all scenarios. Surprisingly, the original implementations of SignatureAnalyzer and SigneR were susceptible to noise, yielding high numbers of false-positive signatures (**Figure 3*b***).

Seven of the tools did not provide an automatic approach for selecting the total number of operative signatures in a dataset (**Table 1**). Instead, most of these tools offered methodologies for manually selecting the optimal number of signatures bringing user-dependence and arbitrariness in selecting solutions. EMu, SigFit, SigMiner, SignatureAnalyzer, SigneR, TensorSignatures, and SigProfilerExtractor provided capabilities for automatically selecting the total number of operative signatures. EMu, TensorSignatures, SigneR select the total number of signatures using Bayesian information criterion (BIC)^41^, while SignatureAnalyzer and SigMiner utilize automatic relevance determination (ARD)^34^. SigFit’s selection approach is based on the Elbow method^42^. SigProfilerExtractor leverages a modified version of the NMFk selection approach which was previously tested on more than 55,000 synthetic random matrices with pre-determined latent factors and shown to outperform other model selection approaches^43^. Importantly, our simulations demonstrate that SigProfilerExtractor’s model selection is robust to noise while the implemented BIC and ARD approaches are affected even by low levels of noise (**Figure 3*b***). In addition to outperforming thirteen other tools on simulated datasets, SigProfilerExtractor can reveal additional biological results as demonstrated by identifying three novel signatures from reanalysis of the PCAWG dataset. Importantly, SigProfilerExtractor identifies signature SBS92 (**Figure 4**) which is associated with tobacco smoking in whole-genome sequenced bladder cancers and in whole-genome sequenced microbiopsies from normal bladder urothelium. The strong transcriptional strand bias observed in SBS92 is indicative of an environmental mutagen exposure that damages purines. Tobacco smoke is a complex mixture of at least 60 chemicals^13^, many capable of causing damage on purines. Interestingly, our and other prior analyses of exome sequenced bladder cancers from The Cancer Genome Atlas (TCGA) project^13, 44^ did not reveal SBS92. Reanalysis of the set of TCGA bladder cancer exomes^45^ with SigProfilerExtractor was also unable to detect SBS92 (**Supplementary Data**). We suspect that the lack of SBS92 in the TCGA bladder cancers was due to the use of exome sequencing; note that SBS92 is predominately found in intergenic regions (**Figure 4*a***) with most samples expected to have less than 15 mutations from SBS92 in their exomes. To confirm this hypothesis, we downsampled the whole-genome sequenced bladder cancers and the whole-genome sequenced microbiopsies from normal bladder urothelium to exomes. SigProfilerExtractor’s analysis of these downsampled genomes was unable to detect SBS92 confirming that exome sequencing is insufficient to identify signature SBS92 (**Supplementary Data**).

In summary, here we report SigProfilerExtractor – a computational tool for *de novo* extraction of mutational signatures. We demonstrate that SigProfilerExtractor outperforms thirteen other tools by conducting the largest benchmarking of bioinformatics approaches for extracting mutational signatures. Further, we apply SigProfilerExtractor to 2,778 whole-genome sequenced cancers and reveal several novel mutational signatures including a signature putatively attributed to tobacco smoking mutagenesis in bladder cancer and in normal bladder epithelium.

## ONLINE METHODS

### Computational implementation of SigProfilerExtractor and its seven modules

The implementation of SigProfilerExtractor can be separated into seven distinct modules which are packaged together into a single bioinformatics tool. *Module 1* processes the initial input data, which can be provided as either a mutational catalogue containing a set of somatic mutations or a mutational matrix. *Module 2* is responsible for resampling and normalization of the mutational matrix prior to performing nonnegative matrix factorization. *Module 3* performs matrix factorization using nonnegative matrix factorization with multiple replicates. *Module 4* utilizes custom clustering to derive consensus solutions and to perform model selection. *Module 5* decomposes the derived set of *de novo* signatures to a set of previously derived COSMIC signatures. *Module 6* is responsible for calculating the activities of different signatures in individual samples. *Module 7* handles the extensive outputting and plotting of the different analysis performed by SigProfilerExtractor. In principle, each of these modules allows extensive customization. SigProfilerExtractor provides a seamless integration of these seven modules that allows using them in an orchestrated and preconfigured manner with little input from a user.

#### Module 1: Processing of input mutational catalogues or input mutational matrices

SigProfilerExtractor deciphers mutational signatures from a mutational matrix ***M*** with *t* rows and *n* columns; rows represent mutational channels while columns reflect individual cancer samples (**Figure 1*a***). The value of each cell in the matrix, ***M***, corresponds to the number of somatic mutations from a particular mutational channel in each sample. The mutational matrix can be provided as a text file with the first column containing the names of the mutational channels and the first row containing the names of the examined samples. Alternatively, users can provide a mutational catalogue of somatic mutations in a commonly used format (*e.g.,* VCF, MAF, *etc.*) and this mutational catalogue will be internally converted into the appropriate mutational matrix by SigProfilerMatrixGenerator^2^.

#### Module 2: Resampling of the input mutational matrix and normalizing the resampled matrix

SigProfilerExtractor does not factorize the original input matrix. Rather, prior to performing matrix factorization, SigProfilerExtractor performs independent Poisson resampling of the original matrix for each replicate^1^. As such, the matrix factorized in each replicate is never the same for a given value of *k* (**Figure 1*b***). The resampling is performed to ensure that Poisson fluctuations of the matrix do not impact the stability of the factorization results. Additional normalization is performed after resampling to overcome potential skewing of the factorization from any hypermutators. SigProfilerExtractor supports four standard normalization methods^46^: *(i)* Gaussian mixture model (GMM) normalization (default); *(ii)* 100X normalization; *(iii)* log2 normalization; *(iv)* no normalization. *No normalization* does not perform any additional transformation on the Poisson resampled matrix. In *log2 normalization*, the sum of each column in the matrix is derived and logarithm with base 2 is calculated for each of these sums. Each cell in a column of the matrix is multiplied by the log2 of the column-sum and subsequently divided by the original column sum. In *100X normalization*, the sum of each column in the matrix is derived. For each column where the sum exceeds 100 times the number of mutational channels (*i.e.*, 100 times the number of rows in the matrix), each cell in the column is multiplied by the 100 times the number of mutational channels and subsequently divided by the original column sum. This normalization ensures that no sample has a total number of mutations above 100 times the number of mutational channels. *GMM normalization* encompasses a two-step process. The first step derives the normalization cutoff value in a data-driven manner using a Gaussian mixture model (GMM). The second step normalizes the appropriate columns using the derived cutoff value. The first step uses a GMM to separate the samples into two groups based on their total number of mutations; the total number of mutations in a sample reflects the sum of a column in the matrix. The group with larger number of samples is subsequently selected, and the same process is applied iteratively until it converges. Convergence is achieved when the mean of the two groups is separated by no more than four standard deviations of the larger group. A cutoff value is derived as the average value plus two standard deviations from the total number of somatic mutations in the last large group. If the derived cutoff value is below 100 times the number of mutational channels, the cutoff value is adjusted to 100 times the number of mutational channels. For each column where the sum exceeds the derived cutoff value, each cell in the column is multiplied by the cutoff value and subsequently divided by the original column sum. Note that no normalization is performed if the means of the first two groups are not separate by at least four standard deviations. In all cases, columns with a sum of zero, reflecting, genomes without any somatic mutations, are ignored to avoid division by zero.

#### Module 3: Matrix Factorization Using Nonnegative Matrix Factorization with Replicates

By default, SigProfilerExtractor factorizes the matrix ***M*** with different ranks searching for an optimal solution between *k=*1 and *k=*40 mutational signatures. For each value of *k*, by default, the tool performs 500 independent nonnegative matrix factorizations of the normalized Poisson resampled input matrix. Thus, for each value of *k*, SigProfilerExtractor generates 500 distinct factorizations of normalized Poisson resampled matrices resulting into 500 different matrices ***S***, each matrix reflecting the patterns of the *de novo* mutational signatures, and 500 different matrices ***A***, each matrix reflecting the activities of the *de novo* mutational signatures (**Figure 1*b***). To perform each of these factorizations, SigProfilerExtractor utilizes a custom implementation of the multiplicative update algorithm^3^. Specifically, SigProfilerExtractor initializes the ***S*** and ***A*** matrices in the first step of the factorization using either random initial conditions (default) or one of the derivatives of nonnegative double singular vector decomposition^47^.

SigProfilerExtractor provides internal support for minimizing three different objective functions based on: *(i)* generalized Kullback-Leibler updates (default); *(ii)* Euclidean updates; *(iii)* Itakura-Saito updates. By default, the tool performs all factorization using multithreading of central processing units (CPUs) and provides support for factorization using graphics processing units (GPUs) by leveraging PyTorch^48^. In all cases, by default, the implemented minimization performs at least 10,000 iterations (also known as NMF updates or NMF multiplicative update steps) with a maximum of 1,000,000 iterations. By default, the convergence tolerance of the algorithm is set to 10^-15^. Note that SigProfilerExtractor allows configuring all factorization parameters.

#### Module 4: Custom partition clustering and performing model selection

The previously described *Module 3* generates a number of sets with each set containing, by default, 500 different matrices ***S***, where each matrix reflects the patterns of *de novo* mutational signatures for a particular factorization of a normalized Poisson resampled matrix. One set, containing 500 different matrices ***S***, is generated for each of the interrogated total number of operative signatures, *k*, with a default range for *k* between 1 and 40 signatures. For each value of *k*, *Module 4* first performs custom clustering of the ***S*** matrices and, subsequently, applies a modified version of the NMFk model selection approach to select the optimal value of *k*^43^ (**Figure 1*b***). Specifically, for each value of *k*, the clustering is initialized with *k* random centroids. One of the ***S*** matrices is randomly chosen, and its columns matched to the most similar centroids with no two columns assigned to the same cluster. The process is repeated until the columns of all ***S*** matrices in the set are assigned to their respective clusters. SigProfilerExtractor implements the Hungarian algorithm^30^ to pair consensus vectors from two matrices (*i.e.,* cluster centroids and mutational signature from a matrix ***S***); the Hungarian algorithm maximizes the total cosine similarities of all paired vectors between two matrices^30^. After assigning all columns to a cluster, the centroids of each cluster are recalculated by evaluating the average of all columns/vectors in a cluster. This process continues iteratively until the average silhouette coefficient converges (*i.e.,* its value does not change by more than 10^-12^). After convergence for a given value of *k*, the centroids of the clusters are reported as consensus mutational signatures, an overall reconstruction error is calculated for describing the original input matrix, ***M***, and stability is calculated for each signature by computing the silhouette value^49^ of the cluster corresponding to that signature (**Figure 1*b***). The silhouette value of a cluster measures the similarities of the objects assigned to that cluster compared to any other cluster. Silhouette values range from −1.0 to +1.0 with values above zero indicating that, on average, objects have a higher similarity with their own cluster compared to their nearest clusters. Note that signatures with low stability correspond to a lack of uniqueness of the NMF due to Poisson resampling and/or to the potential existence of multiple convergent stationary points in the NMF solution^32^.

Our custom clustering is performed for each of the interrogated total number of operative signatures, *k*, with a default range for *k* between 1 and 40 signatures. After performing clustering, for each value of *k*, one has derived: *(i)* the consensus set of mutational signatures; *(ii)* an overall reconstruction error for describing the original input matrix; and *(iii)* stability value for each of the identified consensus mutational signatures.

SigProfilerExtractor performs a solution selection based on the stability of signatures in a solution and the ability of these signatures to reconstruct the original input matrix. By default, SigProfilerExtractor will consider solutions stable if the signatures derived in the solution have an average stability above 0.80 with no individual signature having stability below 0.20. To reduce overfitting, the tool also measures the information gained from the extracted set of signatures in each solution. SigProfilerExtractor compares, using Wilcoxon rank-sum tests, the reconstruction errors across all samples from the stable solution with the greatest number of signatures to the reconstruction errors across all samples from stable solutions with lower number of signatures. Stable solutions with lower number of signatures are compared in a decreasing order to their total number of signatures with comparison stopping if the Wilcoxon rank-sum test yields a *p-value* below 0.05 (*i.e.*, reflecting that a solution does not describe the original data as good as the stable solution with the greatest number of signatures). The stable solution with lowest number of signatures and a Wilcoxon rank-sum test *p-value* above 0.05 is selected as the optimal solution. If no solution has a Wilcoxon rank-sum test *p-value* above 0.05, the stable solution with the greatest number of signatures is selected as the optimal solution. Note that while SigProfilerExtractor selects an optimal solution, it outputs all the information necessary to evaluate mutational signatures and their activities for all other stable and unstable solutions.

#### Module 5: Decomposing de novo extracted signatures to known COSMIC signatures

SigProfilerExtractor provides a module for decomposing each of the *de novo* extracted mutational signatures to a set of previously derived signatures. By default, the tool decomposes each of the signatures in the optimal solution to a set of COSMICv3 reference signatures^14^ with support for signatures of single base substitutions (SBS), doublet base substitutions (DBS), and small insertions and deletions (ID). Since the SBS COSMICv3 reference signatures were derived under the SBS-96 classification^2^, any extended classification of single base substitutions (*e.g.*, SBS-288 and SBS-1536)^2^ is first collapsed to the SBS-96 classification and, subsequently, decomposed to the COSMICv3 reference signatures^14^. The decomposition functionality leverages nonnegative least square (NNLS) algorithm^50^ as its main computational engine. A mixture of addition and removal steps (add-remove functionality) were developed to estimate the list of COSMIC signatures for a *de novo* signature. Specifically, for each *de novo* signature, a COSMIC signature is iteratively added to a list of signatures used to explain the *de novo* signature, NNLS is applied, and the signature which addition causes the greatest decrease of the L2 error is selected. If this greatest decrease is more than a specific threshold (default value of 0.05) then the signature is included in the list of signatures used to explain the *de novo* signature. The addition is immediately followed by a removal step. Each COSMIC signature in the list of signatures used to explain the *de novo* signature are iteratively removed, NNLS is applied, and the signature that causes the least decrease of the L2 error is selected. If this least decrease is less than a specific threshold (default value of 1%) then the signature is removed from the list of signatures used to explain the *de novo* signature. The addition and removal steps are iterated until no signatures are added or removed from the list of signatures used to explain the *de novo* signature. Several previously implemented rules for mutational signatures are incorporated by default in the decomposition module^14^. Specifically, for signatures of single base substitutions: *(i)* the list of signatures used to explain the *de novo* signature is initialized with clock-like signatures SBS1 and SBS5;^11^ *(ii)* biologically connected signatures are included as previously done in Ref ^14^ (*e.g.,* if SBS17a is included in the list then SBS17b is also included the list). The decomposition module is highly customizable as it allows changing all default parameters as well as adding additional new rules or removing existing rules for inclusion and exclusion of particular signatures.

#### Module 6: Evaluating activities of mutational signatures in individual samples

*De novo* extracted and COSMIC derived signatures are refitted to individual samples using nonnegative least squares (NNLS)^50^. *Module 6* internally utilizes the add-remove functionality of *Module 5* with each sample in the original matrix, ***M***, being individually examined. For *de novo* mutational signatures, all *de novo* signatures are initially added to the list of signatures used to explain the sample and a removal step with a cutoff of 2% is applied. To assign COSMIC signatures in a sample, the module first derives the set of *de novo* signatures in that sample. Decomposition to the COSMICv3 signatures using *Module 5* is performed for each of the *de novo* signatures and the identified COSMICv3 signatures are refitted using the add-remove functionality with a removal and addition cutoffs set at 5%. Finally, the activity matrix is constructed by combining the activity vectors generated for all samples in the dataset.

#### Module 7: Outputting and plotting of analysis results

All previous modules make use of *Module 7* for outputting and plotting of the generated results. It should be noted that SigProfilerExtractor provides extensive output for the interrogated total number of operative signatures, *k*, with a default range of *k* between 1 and 40 signatures. For each value of *k*, SigProfilerExtractor outputs the set of operative *de novo* mutational signatures, the activities of the operative signatures, and an extensive set of information related to individual samples, individual *de novo* signatures, and the overall convergence of the factorization and clustering. *Module 7* also provides additional information when ran in debug mode. In addition to outputting information, SigProfilerExtractor also generates a bouquet of plots both for each value of *k* as well as for the suggested optimal solution. SigProfilerExtractor utilizes all previously implemented plots in SigProfilerPlotting^2^ as well as includes several newly developed plots.

### Analysis of the genomics data from cancer and normal somatic tissues

For all examined cancer and normal somatic tissues, *de novo* extraction of mutational signatures was performed with SigProfilerExtractor with default parameters using two distinct mutational classifications: SBS-96 and SBS-288. The SBS-96 mutation classification incorporates the six types of single base substitutions: C>A, C>G, C>T, T>A, T>C, and T>G. Each type of single base substitution is further separated into 16 subtypes determined by the four possible bases 5’ and 3’ adjacent to each mutated base. The SBS-288 mutation classification extends the SBS-96 mutation classification by adding additional information for each of the 96 subtypes. Specifically, SBS-288 incorporates whether a single base substitution is in non-transcribed/intergenic DNA, on the transcribed strand of a gene, or on the untranscribed strand of the gene. *De novo* extraction was performed separately for all examined datasets. Specifically, SigProfilerExtractor was applied: *(i)* to all 2,778 whole-genome sequenced cancers from the Pan-Cancer Analysis of Whole Genomes project^28^; *(ii)* to all samples in each of the 37 cancer types of Pan-Cancer Analysis of Whole Genomes project^28^ with each cancer type examined separately; *(iii)* to all 88 whole-genome sequenced microbiopsies of histologically normal urothelium^35^; *(iv)* to the complete set of bladder cancers from TCGA^45^; *(v)* to exome downsampling of all bladder whole-genome sequenced cancers from the Pan-Cancer Analysis of Whole Genomes project^28^; *(vi)* to exome downsampling of all 88 whole-genome sequenced microbiopsies of histologically normal urothelium^35^. In all cases, the mutational catalogue of each sample was taken from the respective original publications. The results from all performed *de novo* extractions can be found in **Supplementary Data**. Downsampling of a whole-genome sequenced sample to a whole-exome was performed using SigProfilerMatrixGenerator^2^.

### Additional approaches for miscellaneous analysis

Synthetic scenarios were labeled as easy, medium, and hard based on the number of operative signatures in each scenario. Based on our most recent analysis of mutational signatures in 82 cancer types^14^, approximately 7.4% of human cancer types have 5 or less signatures (reflected in simulations of easy scenarios), 15.9% have 11 to 21 signatures (medium scenarios), and 59.5% have 25 or more signatures (hard scenarios). Note that 17.2% of cancer types have either between 5 and 10 signatures or between 22 and 24 signatures.

Cosine similarity was used to compare the profiles of different mutational signatures. P-values can be attributed to cosine similarities based on a null hypothesis of uniform random distribution of nonnegative vectors^33^.

Briefly, the prevalence of somatic mutations in a whole-exome sample was calculated based on the identified mutations in protein coding genes and assuming that an average whole-exome has sufficient coverage of 30.0 megabase-pairs in protein coding genes. The prevalence of somatic mutations in a whole-genome sample was calculated based on all identified mutations and assuming that an average whole-genome has sufficient coverage of 3.00 gigabase-pairs.

All methods related to the generation of the benchmarking scenarios and the application of the different tools to these scenarios can be found in **Supplementary Note 1**.

## SUPPLEMENTARY INFORMATION

**Supplementary Figure 1. Standard set of performance metrics used for benchmarking all bioinformatics tools.** An example demonstrating the derivation of *true positive* (TP), *false positive* (FP), or *false negative* (FN) signatures for a tool applied to a synthetic dataset generated using 6 ground truth signatures (termed, Ground Truth Signatures 1 through 6). The tool extracts 4 signatures (termed, Extracted Signatures A through D). In this example, an extracted signature is considered a true positive if it matches one of the ground-truth signatures with a cosine similarity threshold of at least 0.90.

**Supplementary Table 1. Detailed performance metrics after applying each tool across all SBS-96 noiseless synthetic scenarios.** Performance metrics are calculated as per *Supplementary Figure 1*. An extracted signature is considered a true positive if it matches one of the ground-truth signatures with a cosine similarity threshold of at least 0.90.

**Supplementary Table 2. Detailed performance metrics after applying the seven best performing tools across SBS-96 synthetic scenarios with different levels of noise.** Performance metrics are calculated as per *Supplementary Figure 1*. An extracted signature is considered a true positive if it matches one of the ground-truth signatures with a cosine similarity threshold of at least 0.90.

**Supplementary Table 3. Detailed performance metrics of applying SigProfilerExtractor and SignatureAnalyzer to extended synthetic scenarios.** Performance metrics are calculated as per *Supplementary Figure 1*. An extracted signature is considered a true positive if it matches one of the ground-truth signatures with a cosine similarity threshold of at least 0.90.

**Supplementary Table 4. Profiles of three novel mutational signatures identified in the PCAWG cohort of 2,778 whole-genome sequenced cancers.** The profiles of the novel mutational signatures are reported using the SBS-288 classification which incorporates the trinucleotide context and strand information (intergenic region, untranscribed strand, or transcribed strand) for each type of single base substitution. The SBS-288 classification can be easily collapsed to the commonly used SBS-96 classification.

**Supplementary Note 1. Detailed description of the performed benchmarking.** The supplementary note provides extensive details about each of the generated synthetic scenarios as well as about applying each of the tools to these scenarios. The results from applying all tools to all scenarios, including appropriate input and out files, can be found in **Supplementary Data**.

**Supplementary Data**

All results from the benchmarking with synthetic datasets, including the appropriate input used to run each of the tools as well as the output generated by each of the tools, can be found at: ftp://alexandrovlab-ftp.ucsd.edu/pub/publications/Islam_et_al_SigProfilerExtractor/Benchmark/.

All results from the *de novo* extraction of mutational signatures from the PCAWG dataset can be found at: ftp://alexandrovlab-ftp.ucsd.edu/pub/publications/Islam_et_al_SigProfilerExtractor/PCAWG_Reanalysis/.

All results from the *de novo* extraction of mutational signatures for confirming the patterns of the novel signatures for additional datasets can be found at: ftp://alexandrovlab-ftp.ucsd.edu/pub/publications/Islam_et_al_SigProfilerExtractor/Confirmation_of_Novel_Signatures/.

All results from the *de novo* extraction of mutational signatures from downsampling of whole-genome sequenced samples to whole-exomes can be found at: ftp://alexandrovlab-ftp.ucsd.edu/pub/publications/Islam_et_al_SigProfilerExtractor/Downsampling_of_whole_genomes/

## ACKNOWLEDGEMENTS

The authors would like to thank Allan Balmain (UC San Francisco) for the many useful discussions as well as Ville Mustonen (University of Helsinki) and Israel Tojal Da Silva (A.C. Camargo Cancer Center) for help in configuring EMu and SigneR, respectively. This work was supported by Cancer Research UK Grand Challenge Award C98/A24032 (LBA, PB, and MRS), Wellcome grant reference 206194 (MRS), as well as US National Institute of Health grants R01MH116281-01A1 (BSA), R01ES030993-01A1 (LBA), and R01ES032547 (LBA). This work was also supported by Singapore National Medical Research Council grants NMRC/CIRG/1422/2015 and MOH-000032/MOHCIRG18may-0004 and the Singapore Ministry of Health via the Duke-NUS Signature Research Programmes. LBA is an Abeloff V Scholar and he is supported by an Alfred P. Sloan Research Fellowship. Research at UC San Diego was also supported by a Packard Fellowship for Science and Engineering to LBA. AJG was funded by a postdoctoral fellowship (grant nr. P2BSP3_178591). NP receives funding through the Cancer Research UK Clinician Scientist Fellowship scheme and is supported by University College London Cancer Institute. Research at Los Alamos National Laboratory was conducted under Contract No. 89233218CNA000001 by the U.S. Department of Energy’s National Nuclear Security Administration and supported by Laboratory Directed Research and Development (LDRD) grant 20190020DR (BSA). CDS is supported by the GEM consortium and acknowledges funding for this work through a Cancer Research UK travel grant. The funders had no roles in study design, data collection and analysis, decision to publish, or preparation of the manuscript.

## AUTHOR CONTRIBUTIONS

LBA and SMAI designed both SigProfilerExtractor’s methodology and the performed analyses with help from NP, JZ, DJA, IM, BSA, LH, DCW, MTL, PB, MRS, and SGR. SMAI developed SigProfilerExtractor with help from MV, ENB, YH, CDS, RV, and JW. All synthetic benchmarking datasets were generated by YW and SGR. SMAI documented SigProfilerExtractor and performed the benchmarking of all tools on synthetic data with help from MDG, MB, BO, AK, and AA. Additional validations, confirmations, and applications of SigProfilerExtractor to real and synthetic datasets were performed by SMAI, SM, SS, YRL, NS, LR, TZ, AJG, YH, CDS, and SWB. LBA directed the overall research and wrote the manuscript with help from SMAI and input from all other authors. All authors read and approved the final manuscript.

## TOOL AVAILABILITY

SigProfilerExtractor and all its modules are open source and freely available for use under the permissive 2-clause BSD license. SigProfilerExtractor and its modules are implemented in Python with an R wrapper package allowing users to run the tool from an R environment. SigProfilerExtractor can be installed using the PyPI package manager from https://pypi.org/project/SigProfilerExtractor/ or downloaded from GitHub from https://github.com/AlexandrovLab/SigProfilerExtractor. The R version of the tool can be downloaded from https://github.com/AlexandrovLab/SigProfilerExtractorR. A detailed wiki page including installation, usage, and explanation of result is provided at https://osf.io/t6j7u/wiki/home/. SigProfilerExtractor is compatible with Windows, Linux, Unix, and macOS operating systems.

## REFERENCE

1. Alexandrov, L. B., Nik-Zainal, S., Wedge, D. C., Campbell, P. J. & Stratton, M. R. Deciphering signatures of mutational processes operative in human cancer. Cell Rep 3, 246–259, doi:10.1016/j.celrep.2012.12.008 (2013).

2. Bergstrom, E. N. et al. SigProfilerMatrixGenerator: a tool for visualizing and exploring patterns of small mutational events. BMC Genomics 20, 685, doi:10.1186/s12864-019-6041-2 (2019).

3. Lee, D. D. & Seung, H. S. Learning the parts of objects by non-negative matrix factorization. Nature 401, 788–791, doi:10.1038/44565 (1999).

4. Févotte, C. & Cemgil, A. T. in 2009 17th European Signal Processing Conference. 1913–1917.

5. Dempster, A. P., Laird, N. M. & Rubin, D. B. Maximum Likelihood from Incomplete Data Via the EM Algorithm. Journal of the Royal Statistical Society: Series B (Methodological) 39, 1–22, doi:10.1111/j.2517-6161.1977.tb01600.x (1977).

6. Suri, P. & Roy, N. R. in 2017 3rd International Conference on Computational Intelligence & Communication Technology (CICT). 1–5.

7. Alexandrov, L. B. Understanding the origins of human cancer. Science 350, 1175, doi:10.1126/science.aad7363 (2015).

8. Alexandrov, L. B. et al. Signatures of mutational processes in human cancer. Nature 500, 415–421, doi:10.1038/nature12477 (2013).

9. Petljak, M. & Alexandrov, L. B. Understanding mutagenesis through delineation of mutational signatures in human cancer. Carcinogenesis 37, 531–540, doi:10.1093/carcin/bgw055 (2016).

10. Pich, O. et al. The mutational footprints of cancer therapies. Nat Genet 51, 1732–1740, doi:10.1038/s41588-019-0525-5 (2019).

11. Alexandrov, L. B. et al. Clock-like mutational processes in human somatic cells. Nat Genet 47, 1402–1407, doi:10.1038/ng.3441 (2015).

12. Alexandrov, L. B., Nik-Zainal, S., Siu, H. C., Leung, S. Y. & Stratton, M. R. A mutational signature in gastric cancer suggests therapeutic strategies. Nat Commun 6, 8683, doi:10.1038/ncomms9683 (2015).

13. Alexandrov, L. B. et al. Mutational signatures associated with tobacco smoking in human cancer. Science 354, 618–622, doi:10.1126/science.aag0299 (2016).

14. Alexandrov, L. B. et al. The repertoire of mutational signatures in human cancer. Nature 578, 94–101, doi:10.1038/s41586-020-1943-3 (2020).

15. Fischer, A., Illingworth, C. J., Campbell, P. J. & Mustonen, V. EMu: probabilistic inference of mutational processes and their localization in the cancer genome. Genome Biol 14, R39, doi:10.1186/gb-2013-14-4-r39 (2013).

16. Mayakonda, A., Lin, D. C., Assenov, Y., Plass, C. & Koeffler, H. P. Maftools: efficient and comprehensive analysis of somatic variants in cancer. Genome Res 28, 1747–1756, doi:10.1101/gr.239244.118 (2018).

17. Blokzijl, F., Janssen, R., van Boxtel, R. & Cuppen, E. MutationalPatterns: comprehensive genome-wide analysis of mutational processes. Genome Med 10, 33, doi:10.1186/s13073-018-0539-0 (2018).

18. Fantini, D., Vidimar, V., Yu, Y., Condello, S. & Meeks, J. MutSignatures: an R package for extraction and analysis of cancer mutational signatures. Scientific Reports 10, 18217–18217 (2020).

19. Ardin, M. et al. MutSpec: a Galaxy toolbox for streamlined analyses of somatic mutation spectra in human and mouse cancer genomes. BMC Bioinformatics 17, 170, doi:10.1186/s12859-016-1011-z (2016).

20. Gori, K. & Baez-Ortega, A. sigfit: flexible Bayesian inference of mutational signatures. bioRxiv, 372896, doi:10.1101/372896 (2020).

21. Wang, S. et al. Copy number signature analyses in prostate cancer reveal distinct etiologies and clinical outcomes. medRxiv, 2020.2004.2027.20082404, doi:10.1101/2020.04.27.20082404 (2020).

22. Kasar, S. et al. Whole-genome sequencing reveals activation-induced cytidine deaminase signatures during indolent chronic lymphocytic leukaemia evolution. Nat Commun 6, 8866, doi:10.1038/ncomms9866 (2015).

23. Taylor-Weiner, A. et al. Scaling computational genomics to millions of individuals with GPUs. Genome Biol 20, 228, doi:10.1186/s13059-019-1836-7 (2019).

24. Degasperi, A. et al. A practical framework and online tool for mutational signature analyses show inter-tissue variation and driver dependencies. Nat Cancer 1, 249–263, doi:10.1038/s43018-020-0027-5 (2020).

25. Rosales, R. A., Drummond, R. D., Valieris, R., Dias-Neto, E. & da Silva, I. T. signeR: an empirical Bayesian approach to mutational signature discovery. Bioinformatics 33, 8–16, doi:10.1093/bioinformatics/btw572 (2017).

26. Gehring, J. S., Fischer, B., Lawrence, M. & Huber, W. SomaticSignatures: inferring mutational signatures from single-nucleotide variants. Bioinformatics 31, 3673–3675, doi:10.1093/bioinformatics/btv408 (2015).

27. Vöhringer, H. & Gerstung, M. Learning mutational signatures and their multidimensional genomic properties with TensorSignatures. bioRxiv, 850453 (2019).

28. Consortium, I. T. P.-C. A. o. W. G. Pan-cancer analysis of whole genomes. Nature 578, 82–93, doi:10.1038/s41586-020-1969-6 (2020).

29. Kullback, S. & Leibler, R. A. On Information and Sufficiency. Ann. Math. Statist. 22, 79–86, doi:10.1214/aoms/1177729694 (1951).

30. Kuhn, H. W. The Hungarian method for the assignment problem. Naval research logistics quarterly 2, 83–97 (1955).

31. Huang, K., Sidiropoulos, N. D. & Swami, A. Non-Negative Matrix Factorization Revisited: Uniqueness and Algorithm for Symmetric Decomposition. IEEE Transactions on Signal Processing 62, 211–224, doi:10.1109/TSP.2013.2285514 (2014).

32. Lin, C. On the Convergence of Multiplicative Update Algorithms for Nonnegative Matrix Factorization. IEEE Transactions on Neural Networks 18, 1589–1596, doi:10.1109/TNN.2007.895831 (2007).

33. Bergstrom, E. N., Barnes, M., Martincorena, I. & Alexandrov, L. B. Generating realistic null hypothesis of cancer mutational landscapes using SigProfilerSimulator. BMC Bioinformatics 21, 438, doi:10.1186/s12859-020-03772-3 (2020).

34. Tan, V. Y. F. & Févotte, C. Automatic Relevance Determination in Nonnegative Matrix Factorization with the /spl beta/-Divergence. IEEE Transactions on Pattern Analysis and Machine Intelligence 35, 1592–1605, doi:10.1109/TPAMI.2012.240 (2013).

35. Lawson, A. R. J. et al. Extensive heterogeneity in somatic mutation and selection in the human bladder. Science 370, 75–82, doi:10.1126/science.aba8347 (2020).

36. Moody, S. et al. Mutational signatures in esophageal squamous cell carcinoma from eight countries of varying incidence. medRxiv, 2021.2004.2029.21255920, doi:10.1101/2021.04.29.21255920 (2021).

37. Gaujoux, R. & Seoighe, C. A flexible R package for nonnegative matrix factorization. BMC Bioinformatics 11, 367, doi:10.1186/1471-2105-11-367 (2010).

38. Brunet, J. P., Tamayo, P., Golub, T. R. & Mesirov, J. P. Metagenes and molecular pattern discovery using matrix factorization. Proc Natl Acad Sci U S A 101, 4164–4169, doi:10.1073/pnas.0308531101 (2004).

39. Abadi, M., et al. TensorFlow: Large-Scale Machine Learning on Heterogeneous Distributed Systems. arXiv e-prints, arXiv:1603.04467 (2016).

40. Carpenter, B. et al. Stan: A Probabilistic Programming Language. Journal of Statistical Software 76, doi:10.18637/jss.v076.i01 (2017).

41. Schwarz, G. Estimating the Dimension of a Model. The Annals of Statistics 6, 461–464, doi:10.1214/aos/1176344136 (1978).

42. Thorndike, R. L. Who belongs in the family? Psychometrika 18, 267–276, doi:10.1007/bf02289263 (1953).

43. Benjamin, N., Raviteja, V., Miguel, A. H.-H., Svetlana, K. & Boian, A. A neural network for determination of latent dimensionality in Nonnegative Matrix Factorization. Machine Learning: Science and Technology (2020).

44. Kim, J. et al. Somatic ERCC2 mutations are associated with a distinct genomic signature in urothelial tumors. Nat Genet 48, 600–606, doi:10.1038/ng.3557 (2016).

45. Cancer Genome Atlas Research, N. Comprehensive molecular characterization of urothelial bladder carcinoma. Nature 507, 315–322, doi:10.1038/nature12965 (2014).

46. Shalabi, L. A. & Shaaban, Z. in 2006 International Conference on Dependability of Computer Systems. 207–214.

47. Žitnik, M. & Zupan, B. Nimfa: A python library for nonnegative matrix factorization. The Journal of Machine Learning Research 13, 849–853 (2012).

48. Lew, J. et al. in 2019 IEEE International Symposium on Performance Analysis of Systems and Software (ISPASS). 151–152.

49. Aranganayagi, S. & Thangavel, K. in International Conference on Computational Intelligence and Multimedia Applications (ICCIMA 2007). 13–17 (IEEE).

50. Franc, V., Hlaváč, V. & Navara, M. in Computer Analysis of Images and Patterns. (eds André Gagalowicz & Wilfried Philips) 407–414 (Springer Berlin Heidelberg).

51. Stacklies, W., Redestig, H., Scholz, M., Walther, D. & Selbig, J. pcaMethods--a bioconductor package providing PCA methods for incomplete data. Bioinformatics 23, 1164–1167, doi:10.1093/bioinformatics/btm069 (2007).

