## Supplementary Figure 1 for "Uncovering novel mutational signatures by *de novo* extraction with SigProfilerExtractor"

Simulated *dataset* using 6 ground truth signatures (Ground Truth Signatures 1 through 6). A tool extracts 4 signatures (Extracted Signatures A through D). Comparison between Ground Truth and Extracted Signatures using cosine similarity.

| | Extracted Signature A | Extracted Signature B | Extracted Signature C | Extracted Signature D |
|---|---|---|---|---|
| Ground Truth Signature 1 | 0.14 | **0.98** | 0.56 | 0.36 |
| Ground Truth Signature 2 | 0.35 | 0.29 | **0.93** | 0.46 |
| Ground Truth Signature 3 | 0.31 | 0.56 | 0.78 | 0.66 |
| Ground Truth Signature 4 | 0.34 | 0.08 | 0.57 | 0.67 |
| Ground Truth Signature 5 | **0.95** | 0.15 | 0.81 | 0.39 |
| Ground Truth Signature 6 | 0.23 | 0.74 | 0.48 | 0.26 |

**True Positives (TP; ≥ 0.90)**

Extracted Signature A
Extracted Signature B
Extracted Signature C

Signatures correctly extracted from the *dataset*

**False Positives (FP)**

Extracted Signature D

Signatures extracted but absent in the *dataset*

**False Negatives (FN)**

Ground Truth Signature 3
Ground Truth Signature 4
Ground Truth Signature 6

Signatures not extracted but used in simulating the *dataset*

Cosine similarity between Extracted Signature C and Ground Truth Signature 6

Supplementary Fig. 1
