## Supplementary Note 1 for "Uncovering novel mutational signatures by *de novo* extraction with SigProfilerExtractor"

**Supplementary Note 1. Detailed description of the performed benchmarking.** The supplementary note provides extensive details about each of the generated synthetic scenarios as well as about applying each of the tools to these scenarios. The results from applying all tools to all scenarios, including appropriate input and out files, can be found in **Supplementary Data**.

### Creation of Scenarios with Synthetic Dataset

Benchmarking was performed on simulated datasets with and without noise using the method originally previously described[1]. All datasets without noise are categorized as different scenarios with many of these scenarios attempting to emulate a particular set of cancer types. Specifically, 20 scenarios were created for the SBS-96 mutational classification and 12 additional scenarios were generated for the extended number of channels. For the SBS-96 classification, COSMIC signatures originally extracted from the PCAWG datasets[1] as well as signature extracted using SignatureAnalyzer[1] and random signatures were used as ground-truth signatures. Many of the scenario was created using a combination of tissue specific signatures. Signature profiles of extended scenarios (E) were based either on random signatures or on composite signatures extracted by SignatureAnalyzer. SignatureAnalyzer's composite signatures consist of total of 1,697 mutation types, encompassing an amalgamation of: 1536 strand-agnostic single base substitutions (SBS-1536) in pentanucleotide context, 78 doublet base (DBS-78) substitutions and 83 types of small insertions and deletions (ID-83).

Attributions of signatures in the different scenarios associated with a cancer type, *t*, were generated based on three parameters that were in turn based on the observed statistics for each signature, *s*, in cancer type *t*:

$\pi$, the proportion of tumors of cancer type $t$ with signature $s$

$\mu$, the mean of $\log_{10}$ of the number of $s$ mutations across those tumors of type $t$ that have signature $s$

$\sigma$, the standard deviation of $\log_{10}$ of the numbers of $s$ mutations across those $t$ tumors that have $s$

Detailed description of each of the used scenarios for benchmarking is provided below. Note that some of the generated scenarios were initially created as part of Ref. [1]. The computational approach for generating the synthetic data can be found at: https://github.com/steverozen/SynSigGen.

***Scenarios 1, 2, E-1, and E-2:*** The scenarios were generated to emulate a subset of the pancreatic adenocarcinoma PCAWG dataset with a total 1,000 synthetic samples. Ground-truth signatures were based on COSMIC.

***Scenarios 3, 4, E-3, and E-4:*** Mutational spectra generated from combinations of flat, relatively featureless mutational signatures. A total of 1,000 synthetic tumors emulating a mixture of 500 synthetic Kidney-RCCs (high prevalence and mutation load from SBS5 and SBS40 signatures) and 500 synthetic ovarian adenocarcinomas (high prevalence of and mutation load from SBS3). Ground-truth signatures were based on COSMIC. This dataset embodies tumors with high prevalence of the main flat signatures, SBS3, SBS5, and SBS40, in a realistic context.

***<u>Scenarios 5, 6, E-5, and E-6:</u>*** Mutational spectra generated from signatures with overlapping and potentially interfering profiles. A total of 1,000 synthetic tumors composed from SBS2 and SBS7a, SBS7b. Mutational load distributions were drawn from bladder transitional cell carcinoma (SBS2) and skin melanoma (SBS7a, SBS7b). Ground-truth signatures were based on COSMIC. Most spectra contain both signatures. The potential interference is between SBS2 (mainly C>T) and SBS7a, SBS7b (mainly C>T).

***<u>Scenarios 7, 8, E-7, and E-8</u>:*** Mutational spectra generated from combinations of flat, relatively featureless mutational signatures. A total of 1,000 synthetic tumors emulating a mixture of 500 synthetic Kidney-RCCs (high prevalence and mutation load from SBS5 and SBS40 signatures) and 500 synthetic ovarian adenocarcinomas (high prevalence of and mutation load from SBS3). Ground-truth signatures were based on signatures extracted by SignatureAnalyzer. This dataset embodies tumors with high prevalence of the main flat signatures, SBS3, SBS5, and SBS40, in a realistic context.

***<u>Scenarios 9, 10, E-9, and E-10:</u>*** Mutational spectra generated from signatures with overlapping and potentially interfering profiles. A total of 1,000 synthetic tumors composed from SBS2 and SBS7a, SBS7b. Mutational load distributions were drawn from bladder transitional cell carcinoma (SBS2) and skin melanoma (SBS7a, SBS7b). Ground-truth signatures were based on signatures extracted by SignatureAnalyzer. Most spectra contain both signatures. The potential interference is between SBS2 (mainly C>T) and SBS7a, SBS7b (mainly C>T).

***Scenarios 11, 12, E-11, and E-12:*** A set of 30 random mutational signature profiles based on SBS-96 classification and a set of 30 random 1,697 feature signature profiles (mimicking SignatureAnalyzer's composite signatures). Each of these sets of random signatures were used in two types of exposures, one with more (mean ~15.6) signatures per tumor and one with fewer (mean ~4) signatures per tumor.

***Scenarios 13 and 14:*** A set of 2,700 synthetic whole-genome samples with mutational spectra matching the ones observed in PCAWG, including 300 spectra from each of 9 cancer different types. These spectra consist of 300 synthetic spectra from each of the following cancer types: bladder transitional cell carcinoma, esophageal adenocarcinoma, breast adenocarcinoma, lung squamous cell carcinoma, renal cell carcinoma, ovarian adenocarcinoma, osteosarcoma, cervical adenocarcinoma, and stomach adenocarcinoma. Ground-truth signatures were based on COSMIC as well as on signatures extracted by SignatureAnalyzer.

***Scenarios 15 and 18:*** A set of 5 random mutational signature profiles based on SBS-96 mutational classification. A total of 1,000 synthetic tumors were generated with one scenario containing an average of 3 signatures per tumor while the other scenario had an average of 5 signatures per tumor.

***Scenarios 16 and 19:*** A set of 15 random mutational signature profiles based on SBS-96 mutational classification. A total of 1,000 synthetic tumors were generated with one scenario containing an average of 10 signatures per tumor while the other scenario had an average of 5 signatures per tumor.

***Scenarios 17 and 20:*** A set of 25 random mutational signature profiles based on SBS-96 mutational classification. A total of 1,000 synthetic tumors were generated with one scenario containing an average of 15 signatures per tumor while the other scenario had an average of 5 signatures per tumor.

Scenario 1 to 14 had only a single replicate while scenario 15 through 20 had 10 replicates each. In addition to noiseless scenarios, we simulated 20 replicates of a scenario with noise: 10 of the replicates were based on scenario 11 and another 10 replicates were based on scenario 12. In each case, while Gaussian noise was added to each replicate. Specifically, random noise for each datapoint (*i.e.,* reflecting a number of mutations of a specific mutation type in a cancer sample) was added with mean value corresponding to the datapoint and standard deviation reflecting 0%, 1%, 2.5%, 5%, or 10% of the value of the data point. Overall, 5 distinct levels of noise were generated, each repeated 20 times, with an average noise level corresponding to 0%, 1%, 2.5%, 5%, and 10% of all mutations observed in the replicate.

**<u>Benchmarking bioinformatic tools for *de novo* extraction of mutational signatures</u>**

The hitherto described synthetic scenarios were used to compare SigProfilerExtractor and thirteen other tools for *de novo* extraction of mutational signatures. The method and parameters we used to extract signatures from the simulated dataset using each tool are described below. With the exception of SignatureAnalyzer, which supports only automatic detection of the total number of mutational signatures without a prespecified range, all other tools required specifying the range for the total number of operative mutational signatures. The rage for benchmarking with suggested model selection, which most closely matches analysis of a real dataset with unknown number of signatures, is provided for each of the scenarios. Benchmarking with forced model selection, where tools were required to extract the known number of ground-truth mutational signatures, performed *de novo* extraction based on the ground truth number of total mutational signatures.

| **Scenario(s)** | **Suggested Selection**<br>Range of Signatures | **Forced Selection**<br>Ground-truth number |
|---|---|---|
| **Scenarios 1 and E-1** | 1 to 30 | 20 |
| **Scenarios 2 and E-2** | 1 to 20 | 11 |
| **Scenarios 3 and E-3** | 1 to 30 | 19 |
| **Scenarios 4 and E-4** | 1 to 20 | 11 |
| **Scenarios 5 and E-5** | 1 to 30 | 26 |
| **Scenarios 6 and E-6** | 1 to 20 | 11 |
| **Scenarios 7 and E-7** | 1 to 10 | 3 |
| **Scenarios 8 and E-8** | 1 to 10 | 3 |

| | | |
|---|---|---|
| **Scenarios 9 and E-9** | 1 to 10 | 3 |
| **Scenarios 10 and E-10** | 1 to 10 | 3 |
| **Scenarios 11 and E-11** | 1 to 35 | 30 |
| **Scenarios 12 and E-12** | 1 to 35 | 30 |
| **Scenarios 13** | 1 to 45 | 39 |
| **Scenarios 14** | 1 to 30 | 21 |
| **Scenarios 15** | 1 to 10 | 5 |
| **Scenarios 16** | 1 to 20 | 15 |
| **Scenarios 17** | 1 to 30 | 25 |
| **Scenarios 18** | 1 to 10 | 5 |
| **Scenarios 19** | 1 to 20 | 15 |
| **Scenarios 20** | 1 to 30 | 25 |

***SigProfilerExtractor:*** The default settings of SigProfilerExtractor (version 1.0.19) were used to extract mutational signatures with minor modifications to reduce overall extraction time. Specifically, we utilized "NMF replicates" = 100, "minimum NMF iterations" = 1,000, "maximum NMF iterations" = 200,000, "NMF test convergence" =1,000 and "NMF tolerance" = 1e-08 parameter settings all other scenarios without noise all replicates with noise.

SigProfiler_PCAWG[1]: The default settings of SigProExtractor (version 0.0.5.48) were used to extract signatures from the benchmark scenarios with exception of "totaliteration"=100.

***SignatureAnalyzer*[2,3]*:*** For the scenarios with and without noise, we used the default parameters described in https://github.com/broadinstitute/SignatureAnalyzer-GPU. For the extended scenarios, the CPU version of SignatureAnalyzer was used with 20 runs with default parameters. The mode number of signatures counts were selected for further evaluation.

***MutationalPatterns*[4]*:*** We downloaded MutationalPatterns version 3.0.1 according to the instruction at https://bioconductor.org/packages/release/bioc/html/MutationalPatterns.html. To extract signatures, we used the NMF method with default parameters with the exception of the "nrun" parameter. The "nrun" parameter was set to 200 in order to increase the reliability of the extraction. To select the optimum number of signatures, as suggested by the developers of the tool, we used the RSS plot that is generated in the NMF rank survey plot.

***SigantureToolsLib*[5]*:*** We downloaded SignaturesToolsLib from https://github.com/Nik-Zainal-Group/signature.tools.lib and the tool was used with default parameters. Depending on the size of datasets and the efficiency of the tool, we used "no filter" approach with 100 bootstrap catalogs. As suggested by the developers, we selected the optimum number of signatures from the plot illustrated the overall metrics. We mostly used the "norm.error" and "Ave.SilWid" with Clustering with Matching (MC) to selected the total number of operative mutational signatures.

***SigneR*[6]*:*** We used the SigneR version 1.16.0 as described in http://bioconductor.org/packages/release/bioc/vignettes/signeR/inst/doc/signeR-vignette.html. The signatures were extracted with default parameters without using an opportunity matrix.

***MutSpec*[7]:** We used the command line platform of MutSpec version 2.0, as described at https://github.com/IARCbioinfo/mutspec. To extract signatures, we used the default parameters. As suggested by the developers, we estimated the optimum number of signatures from the "NMF rank survey" plot generated from the "MutSpec-NMF_Estimate_Signatures" module of the package.

***SomaticSignatures*[8]:** We followed the instructions described at https://www.bioconductor.org/packages/release/bioc/vignettes/SomaticSignatures/inst/doc/SomaticSignatures-vignette.html to use the NMF method to access and extract mutational signatures. SomaticSignatures version 2.26.0 was used with default parameters. To access the number of signatures, we increased the value of "nmf_replicates" from 5 to 30 in order to get better reproducibility. As suggested by the developers, we selected the optimum number of signatures using the "plotNumberSignatures" function provided by the tool. In the plots, we relied on the RSS and explained variance value to choose the optimum solution.

***Maftools*[9]:** We followed the instructions provided at https://www.bioconductor.org/packages/release/bioc/vignettes/maftools/inst/doc/maftools.html to download and extract signatures using Maftools version 2.2.0. As suggested by the developers, we estimated the goodness of fit to decide the optimal number of signatures using the "estimateSignatures" function. Then we extracted the corresponding optimal number signatures using the "extractSignatures" function provided by the tool. All settings were kept as defaults, except we increased the value of "nTry" from 6 to 20 to increase reproducibility.

***SigMiner*[10]*:*** We have SigMiner version 1.0.0 according to the instructions provided at https://shixiangwang.github.io/sigminer-doc/. All the parameters were set as defaults to both estimate as well as extract mutational signatures. To select the optimum number of signatures, as suggested by the developers, we assessed the statistics provided in the NMF rank survey plot.

***SigFit*[11]*:*** We used the instructions provided at http://htmlpreview.github.io/?https://github.com/kgori/sigfit/blob/master/doc/sigfit_vignette.html to download and extract signatures from SigFit version 2.0.0. Signature extraction was done using default parameters with the exception of the total number of iterations which was set at 100.

***EMu*[12]*:*** Benchmarking for EMu was done with v.1.5.2 using default parameters for suggested extraction. Additionally, the optional force parameter was used for benchmarks done with a specific number of processes. EMu was the only tool that was unable to complete *de novo* extractions from a number of synthetic datasets (**Supplementary Tables 1-3**) with the tool either running out on instances with 256 GiB memory or running for 4+ weeks without producing any results. These scenarios were considered as failed and assigned $F_1$ scores of zero.

***MutSignatures*[13]*:*** MutSignatures was downloaded and run according to https://cran.r-project.org/web/packages/mutSignatures/vignettes/get_sarted_with_mutSignatures.html. Signatures were extracted using 500 iterations (num_totIterations = 500).

***TensorSignatures[14]:*** TensorSignatures was downloaded and run according to the instructions at https://github.com/sagar87/tensorsignatures. Input VCFs were generated from the matrices by running SigProfilerSimulator[15]. The headers of the VCF files were modified for TensorSignatures to compute the trinucleotide normalization. TensorSignatures was applied using 10,000 epochs, overdispersion of 50, and trinucleotide normalization. Each decomposition rank was run with 10 iterations. TensorSignatures was not applied to scenarios 13 and 14 as the expected computation time, even with multiple GPUs, was expected to be more than 6 months per scenario.

### SUPPLEMENTARY REFERENCE


1. Alexandrov, L.B. *et al.* The repertoire of mutational signatures in human cancer. *Nature* **578**, 94-101 (2020).
2. Kasar, S. *et al.* Whole-genome sequencing reveals activation-induced cytidine deaminase signatures during indolent chronic lymphocytic leukaemia evolution. *Nat Commun* **6**, 8866 (2015).
3. Taylor-Weiner, A. *et al.* Scaling computational genomics to millions of individuals with GPUs. *Genome Biol* **20**, 228 (2019).
4. Blokzijl, F., Janssen, R., van Boxtel, R. & Cuppen, E. MutationalPatterns: comprehensive genome-wide analysis of mutational processes. *Genome Med* **10**, 33 (2018).
5. Degasperi, A. *et al.* A practical framework and online tool for mutational signature analyses show inter-tissue variation and driver dependencies. *Nat Cancer* **1**, 249-263 (2020).
6. Rosales, R.A., Drummond, R.D., Valieris, R., Dias-Neto, E. & da Silva, I.T. signeR: an empirical Bayesian approach to mutational signature discovery. *Bioinformatics* **33**, 8-16 (2017).
7. Ardin, M. *et al.* MutSpec: a Galaxy toolbox for streamlined analyses of somatic mutation spectra in human and mouse cancer genomes. *BMC Bioinformatics* **17**, 170 (2016).
8. Gehring, J.S., Fischer, B., Lawrence, M. & Huber, W. SomaticSignatures: inferring mutational signatures from single-nucleotide variants. *Bioinformatics* **31**, 3673-5 (2015).
9. Mayakonda, A., Lin, D.C., Assenov, Y., Plass, C. & Koeffler, H.P. Maftools: efficient and comprehensive analysis of somatic variants in cancer. *Genome Res* **28**, 1747-1756 (2018).
10. Wang, S. *et al.* Copy number signature analyses in prostate cancer reveal distinct etiologies and clinical outcomes. *medRxiv*, 2020.04.27.20082404 (2020).
11. Gori, K. & Baez-Ortega, A. sigfit: flexible Bayesian inference of mutational signatures. *bioRxiv*, 372896 (2020).
12. Fischer, A., Illingworth, C.J., Campbell, P.J. & Mustonen, V. EMu: probabilistic inference of mutational processes and their localization in the cancer genome. *Genome Biol* **14**, R39 (2013).
13. Fantini, D., Vidimar, V., Yu, Y., Condello, S. & Meeks, J. MutSignatures: an R package for extraction and analysis of cancer mutational signatures. *Scientific Reports* **10**, 18217-18217 (2020).
14. Vöhringer, H. & Gerstung, M. Learning mutational signatures and their multidimensional genomic properties with TensorSignatures. *bioRxiv*, 850453 (2019).
15. Bergstrom, E.N., Barnes, M., Martincorena, I. & Alexandrov, L.B. Generating realistic null hypothesis of cancer mutational landscapes using SigProfilerSimulator. *BMC Bioinformatics* **21**, 438 (2020).
